# A functional study of all 40 *C. elegans* insulin-like peptides

**DOI:** 10.1101/336750

**Authors:** Shanqing Zheng, Hilton Chiu, Jeffrey Boudreau, Tony Papanicolaou, William Bendena, Ian Chin-Sang

## Abstract

The human genome encodes ten insulin-like genes, whereas the *C. elegans* genome remarkably encodes forty insulin-like genes. The roles of insulin/insulin-like peptide ligands (INS) in *C. elegans* are not well understood. The functional redundancy of the forty INS genes makes it challenging to address their functions by using knock out strategies. Here, we individually overexpressed each of the forty *ins* genes pan-neuronally, and monitored multiple phenotypes including: L1 arrest life span, neuroblast divisions under L1 arrest, dauer formation and fat accumulation, as readouts to characterize the functions of each INS *in vivo*. Of the 40 INS peptides, we found functions for 35 INS peptides and functionally categorized each as agonists, antagonists or of pleiotropic function. In particular, we found that 9 of 16 agonistic INS peptides shortened L1 arrest life span and promoted neuroblast divisions during L1 arrest. Our study revealed that a subset of β-class INS peptides that contain a distinct F peptide sequence are agonists. Our work is the first to categorize the structures of INS peptides and relate these structures to the functions of all forty INS peptides *in vivo*. Our findings will promote the study of insulin function on development, metabolism, and aging-related diseases.

**Author Summary:** Insulin and insulin-like growth factors are found in all animals and regulate many physiological and developmental processes. The human genome has 10 insulin-like peptides including the well characterized insulin hormone. The nematode *C. elegans* has 40 insulin-like (INS) peptide genes. All 40 INS peptides have been knocked out but no single INS gene knock out resembles the loss of the *C. elegans* insulin receptor suggesting that the other INS peptides can compensate when one INS is lost. We have used a genetic approach to overexpress each of the 40 INS peptides in *C. elegans* and have identified *in vivo* function for 35 of the 40 INS peptides. Like the human insulin and IGF-1, *C. elegans* INS peptides are derived from a precursor protein and we have shown that INS peptides with an associated peptide called the F peptide are strong activators of the *C. elegans* insulin-like receptor. We also identified several INS peptides that inhibit the insulin-like receptor and these inhibitory INS peptides may have therapeutic potential.

## Introduction

The *C. elegans* insulin/insulin-like growth factor signaling (IIS) pathway has been extensively studied and the IIS pathway components are evolutionary conserved in metazoans [1]. Insulin-like (INS) peptides bind to and activate cell surface receptors with intrinsic tyrosine kinase activity [2,3]. Auto-phosphorylation of the receptors promote the recruitment and activation of downstream components to initiate their biological effects [4]. Unlike the components in the IIS pathway signal transduction, INS peptides in *C. elegans* are not well studied. The insulin superfamily genes are ubiquitous and have been identified in all animals [5,6]. Compared to human and *Drosophila*, which have ten and eight INS peptides respectively, the *C. elegans* genome encodes forty INS genes [5,7], suggesting that there may be more functional diversity among the *C. elegans* INS peptides. Strikingly, there is only one INS receptor, DAF-2/INSR, which the forty INS peptides are thought to bind as ligands. To date, no single loss of function mutation can fully recapitulate the phenotypes associated with loss of the *daf-2* insulin receptor [8]. The lack of loss-of-function phenotypes for many of the INS peptides suggests that some act redundantly in *C. elegans*, and this makes addressing INS functions by knock-out strategies challenging and limited [8].

In mammals (including humans), INS peptides and IIS signaling control glucose levels, hormone homeostasis and metabolism [9]. In *C. elegans*, IIS signaling controls aging, development, behavior, dauer formation, as well as fat accumulation [10]. By using dauer formation as a readout phenotype, only INS-4, 6 and DAF-28 were identified as potential agonists, while INS-1, 17 and 18 were identified as potential antagonists [7,11,12]. Some INS peptides were suggested to be potential agonists or antagonists based on mRNA expression dynamics between fed and starved conditions [13] and the INS peptides have been shown to regulate each other transcriptionally [14]. However, the roles and functions of all forty INS peptides remain unclear. INS peptides are expressed primarily in the nervous system [7]. As such, we individually overexpressed each of the forty INS peptides in all neurons using a pan-neuronal promoter to direct gene expression [15]. Overexpression (*oe)* lines were then characterized based on phenotypes associated with abnormal IIS signaling in *C. elegans*. As an example, in the absence of food, *C. elegans* can arrest development in the first larval stage (L1 arrest), preventing further growth and development [16]. The level of INS signaling can shorten or lengthen the life span of L1 arrested animals. Here, we assayed the contribution of individual INS peptides on phenotypes associated with IIS signaling that include alterations in L1 arrest life span, Q cell neuroblast divisions, dauer formation and fat metabolism. Based on these assays, seven INS peptides (INS-3, 4, 6, 9, 19, 32, DAF-28) were categorized as strong agonists and three INS peptides (INS-17, 37, 39) were strong antagonists of the DAF-2 INS receptor *in vivo*. Nine INS peptides (INS-1, 2, 10, 11, 13, 20, 24, 29, 35) were found to be weak agonists and five INS peptides (INS-15, 21, 22, 36, 38) functioned as weak antagonists. Five INS peptides (INS-5, 23, 26, 27, 33) did not exhibit any significant phenotype *in vivo*. Eleven of the forty INS peptides have diverse roles serving as agonists or antagonists of DAF-2/INSR depending on the phenotypes scored. The forty INS peptides in *C. elegans* have been grouped into three classes: α, β and γ (Figure 7), on the basis of predicted arrangements of their disulfide bonds [7]. Our work revealed that the β class INS peptides that contain a sequence known as the F peptide is a strong predictor of an INS with activation properties. The majority of β class INS function as agonistic ligands of DAF-2/INSR. Our INS overexpression work reveals the functional nature of signaling for each of the forty INS *in vivo*, and promotes future studies on the functions of the entire *C. elegans* insulin gene family on aging, development and metabolic diseases.

## Results

### Overexpressed INS affect L1 arrest life span

In the absence of food, newly hatched *C. elegans* larva (L1 stage) undergo a developmental quiescence called L1 arrest. Previously, we and others have shown that down regulation of the IIS pathway is critical for L1 arrest survival [17,18]. Wild type L1 arrested worms live for a maximum of 20 days with a mean life span of 13 days when grown at 20°C. Manipulation of IIS signaling can alter the normal 20 day survival period. DAF-18 is the worm orthologue of the human PTEN tumor suppressor. DAF-18/PTEN functions to inhibit the IIS pathway. Enhanced IIS caused by the loss of *daf-18* resulted in shortened life span during L1 arrest (Figure 1A). Blocking IIS by loss of the *daf-2*/INSR resulted in lengthened lifespan in L1 arrested worms (Figure 1A). To categorize the functional role of INS peptides on the regulation of the IIS pathway, each of the forty *ins* (*oe*) strains were scored for L1 arrest life span. We found that twenty-one (*ins-1, 2, 3, 4, 6, 7, 8, 9, 10, 11, 13, 16, 18, 19, 20, 25, 29, 30, 31, 32* and *daf-28*) *ins* (*oe*) worms had significantly shorter life span when compared to wild type worms during L1 arrest (Figure 1B and C). This suggests that these twenty-one INS peptides function as DAF-2/INSR agonists to activate the IIS pathway. Eight (*ins-12, 14, 17, 22, 28, 34, 37*, and *39*) *ins* (*oe*) worms had significantly longer life span than wild type L1 worms (Figure 1D), suggesting that these INS function as DAF-2/INSR antagonists to shut down IIS. Eleven *ins* (*oe*) (*ins*-*5, 15, 21, 23, 24, 26, 27, 33, 35, 36* and *38*) strains had normal L1 arrest life span compared to wild type worms (Supplemental sheet), suggesting that these INS peptides play neutral roles on L1 arrest life span.

**Figure 1.**
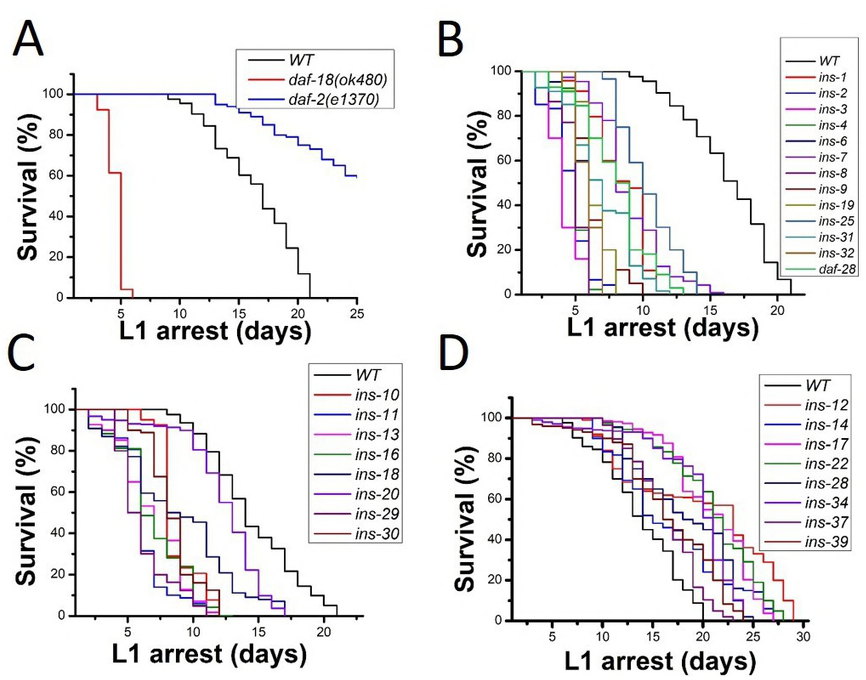
*ins* (*oe*) functions on L1 arrest life span. (A) *daf-18/pten* mutants have shorter L1 arrest life span and the insulin receptor *daf-2* mutants have a longer L1 arrest life span than wild type worms. (B, C) 21 *ins* (*oe*) strains have shorter L1 life span, suggesting these INS are DAF-2 agonists. (D) 8 *ins* (*oe*) strains have longer L1 life span, suggesting these INS peptides are DAF-2 antagonists. Also see details in Supplemental sheet.

### Agonistic INS peptides cause Q cell divisions during L1 arrest

During L1 arrest all cell divisions are halted due, in part, to the shutdown of the IIS pathway. When the IIS pathway is activated during L1 arrest, we showed that the Q cell lineage undergoes cells divisions and movements (in review). We asked whether INS (*oe*) could cause Q cell divisions during L1 arrest. To aid with the scoring of the Q cell divisions in L1 arrest, we scored the presence of the Q cell neuroblast descendants AVM and PVM. Normally, L1 arrested worms only have the embryonic mechanosensory neurons ALMs and PLMs (Figure 2A). Our previous work found that AVM and PVM were present in L1 arrested *daf-18* mutants (Figure 2B), as loss of *daf-18* enhances insulin signaling. Therefore in L1 arrest, if A/PVM are present it tells us the Q cell lineage has undergone its terminal divisions and can be used as a new readout to analyze the functions of the INS ligands.

**Figure 2.**
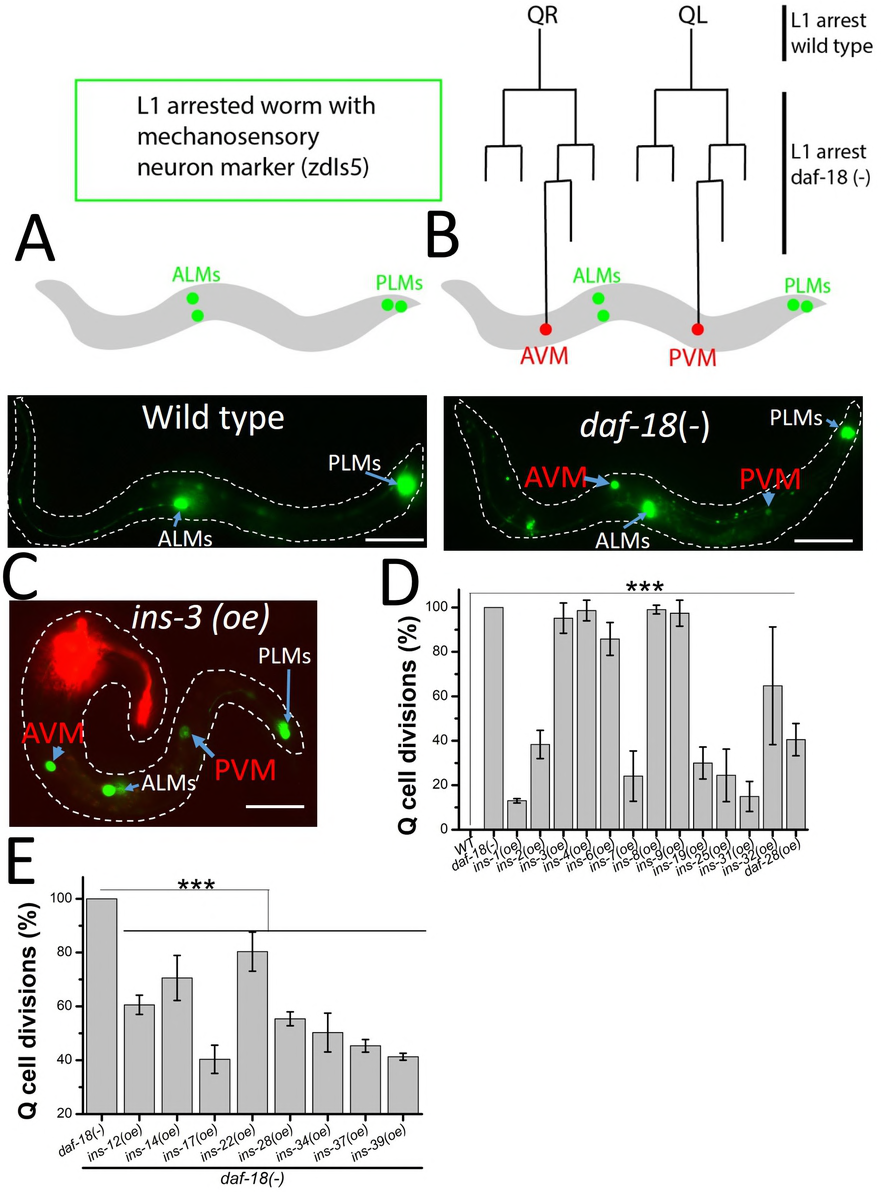
*ins* (*oe*) functions on L1 arrest Q cell divisions. (A) Wild type worms stop development at L1 arrest. The wild type worms with touch neuron marker (*zdIs5*) only have embryonic ALMs and PLMs. (B) *daf-18 (-)* L1 arrest mutants have two terminal Q cell descendants AVM and PVM. (C) A representative *ins-3* (*oe*). The red fluorescence is the AWC neuron from *odr-1∷rfp* transgenic marker for *ins (oe)* lines. (D) 13 *ins* (*oe*) strains show L1 arrest Q cell divisions, suggesting these are DAF-2 agonists. (D) 8 *ins* (*oe*) antagonists which have longer L1 arrest life span (Figure 1D) can suppress the L1 arrest Q cell divisions in *daf-18 (-)* mutants, suggesting these INS are DAF-2 antagonists for Q cell divisions. Scale bars represent 50 µm. Error bars represent the SD. *** P value vs control <0.001. Also see details in Supplemental sheet.

We found that thirteen INS *(oe)* strains (*ins-1*, *2*, *3*, *4*, *6*, *7*, *8*, *9*, *19*, *25*, *31*, *32* and *daf-28)* caused Q cell divisions in L1 arrested worms (Figure 2C and D). Our results suggested that these INS peptides are strong DAF-2 agonists. The Q cell divisions during L1 arrest is an excellent readout for INS peptides that activate the receptor. However, this phenotype on its own is not sufficient to determine INS peptides that are inhibitory to DAF-2 as the loss of DAF-2 is the same as wildtype (i.e. no Q cell divisions during L1 arrest). To examine INS peptides found to be inhibitory based on L1 arrest life span, we used *daf-18* mutant worms (i.e. increased IIS signaling) and asked whether the inhibitory INS peptides could suppress the *daf-18* Q L1 arrest cell divisions. As predicted, we found that all the *ins* (*oe*) strains which could make L1 arrested worms live longer (Figure 1D) could also significantly suppress the *daf-18* Q cell divisions (Figure 2E). These results confirm that the eight INS peptides (INS-12, 14, 17, 22, 28, 34, 37, and 39) function as DAF-2/INSR antagonists in both L1 arrest life span and Q cell divisions. All INS peptides that could induce L1 arrest Q cell divisions also shortened L1 arrest lifespan. However, not all the *ins* genes which shorten L1 arrest lifespan can induce L1 arrest Q cell divisions (Figure 1C), suggesting that these INS peptides have specific activities in controlling L1 arrest life span and cell divisions.

To ensure that INS peptides overexpressed in the nervous system were dependent on normal peptide processing, we tested whether the pro-protein convertase deficient animal (*egl-3* mutant) [12] could suppress the function of agonistic INS peptides on L1 arrest Q cell divisions. We found that L1 arrested Q cell divisions in *ins (oe)* worms were completely suppressed by *egl-3*. In addition, if these agonists activate the DAF-2/INSR then *daf-2* mutants should suppress the *ins (oe)* L1 arrest Q cell divisions. We found that each *ins (oe)* strain that induced L1 arrested Q cell divisions was completely suppressed in the *daf-2(e979)* or *daf-2 (e1370)* mutant background (data not show). Previous work showed that activated IIS pathway also induced germ cell and M cell divisions in L1 arrested worms [13,19], therefore we determined whether these overexpressed INS peptides would enhance germ cell divisions and/or M cell divisions. Indeed, the overexpressed agonistic INS peptides were able to act cell non-autonomously, resulting in M cell and germ cell divisions during L1 arrest (Figure 3).

**Figure 3.**
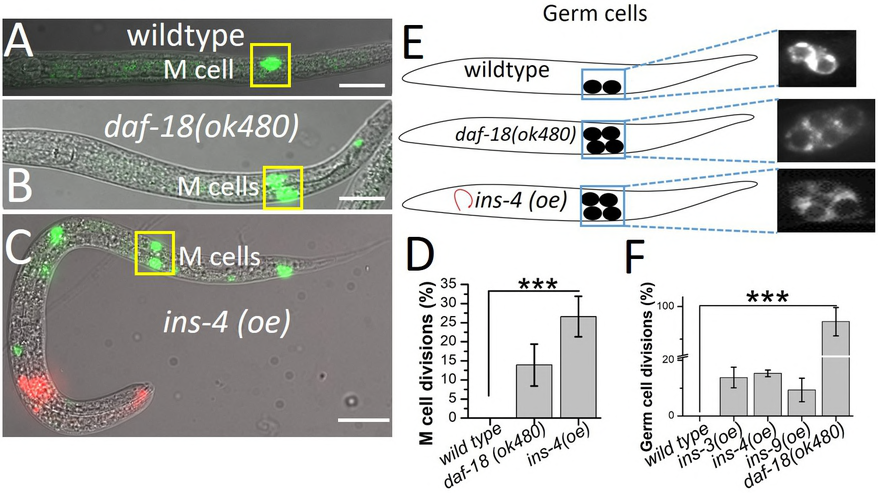
Pan-neuronal INS overexpression acts cell non-autonomously to induce non neuronal cell divisions during L1 arrest. (A) L1 arrest wild-type, only one M cell is observed (*ayIs6*). (B, C, D) *daf-18 (-)* and Pan-neuronal INS overexpression cause the M cell to divide in L1 arrest. (E, F) Germ cells (Z2/Z3) in L1 arrested worms. *daf-18(-)* mutants and Pan-neuronal INS overexpression induces germ cell divisions. Red florescence in the head is the (*odr-1∷rfp*): transgenic marker. Data represents the average of at least 3 independent experiments from at least two stable transgenic lines. Scale bars represent 50 µm. Error bars represent the SD. *** P value vs control <0.001.

### INS peptide function on dauer formation

In *C. elegans*, animals in the second larval stage can enter a dauer diapause phase under adverse environmental conditions. Mutants that reduced IIS (such as *daf-2* INSR (*lf*) or *age-1* PI3K (*lf*)) have a dauer-constitutive (Daf-c) phenotype. Yet, individual INS knock out mutants only show a very weak dauer phenotype which may imply functional redundancy [14]. Functional redundancy was supported by creating a fully penetrant Daf-c phenotype by simultaneous removal of *ins-4, ins-6* and *daf-28* (Hung et al. 2014). However, the functions of the forty *ins* genes on dauer formation are still not well addressed. Here, we tested the function of individual pan-neuronal INS on dauer formation. Previous work showed that wildtype *C. elegans* can go into dauer under high temperatures even in the presence of food or non-crowding conditions [20]. At 29° C we found nineteen INS peptides (INS-1, 2, 3, 4, 6, 8, 9, 10, 11, 12, 13, 14, 19, 24, 28, 32, 34, 35 and DAF-28) significantly reduced dauer formation (Figure 4C), consistent with these INS peptides acting as DAF-2/INSR agonists. Twelve INS peptides (INS-7, 15, 16, 17, 18, 25, 30, 31, 36, 37, 38 and 39) caused higher dauer penetrance than wild type (Figure 4B), consistent with these INS peptides acting as DAF-2/INSR antagonists. Nine INS peptides (INS-5, 20, 21, 22, 23, 26, 27, 29 and 33) had no significant functions on dauer formation (Figure 4A). Interestingly, with ten INS peptides (INS-1, 2, 3, 4, 6, 8, 9, 19, 32 and daf-28) worms showed shortened L1 arrest life span, promoted L1 arrest Q cell divisions and reduced dauer formation, suggesting that these ten INS can activate DAF-2 function consistently when scored by different phenotypes. Notably, not all the INS peptides which extended L1 arrest life span enhanced dauer formation, suggesting that these INS peptides have different functions to control L1 arrest and dauer. For example, INS-12, 14, 22, 28, and 34 acted like DAF-2/INSR antagonists as they increased L1 life span, while they acted like agonists or had no function in dauer formation. INS-7, 16, 18, 25, 30 and 31 acted as DAF-2/INSR agonists as they reduced L1 arrest life span, but acted as DAF-2/INSR antagonists by increasing dauer formation.

**Figure 4.**
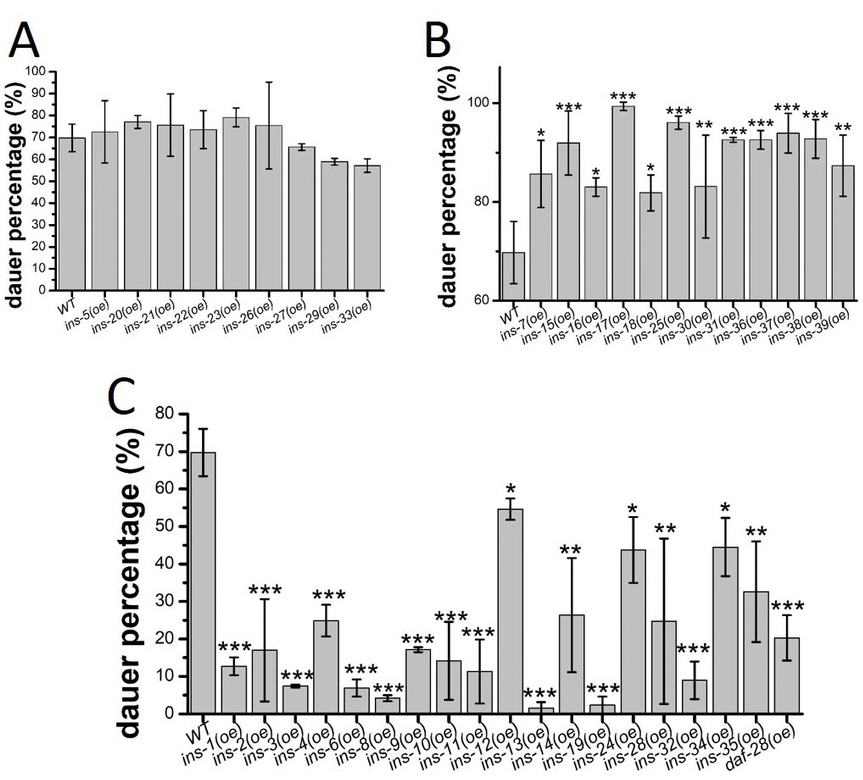
*ins* (*oe*) strains on dauer formation at high temperature. (A) 9 *ins* (*oe*) worms have no significant functions on dauer formation. (B) 12 *ins* (*oe*) worms (antagonists) induce more dauer than wild type worms. (C) 19 *ins* (*oe*) worms (agonists) have less dauer than wild type worms. Error bars represent the SD, * P value vs control <0.05, ** P value vs control <0.01 *** P value vs control <0.001. Also see details in Supplemental sheet.

### INS peptide function on fat accumulation in adult worms

The IIS pathway plays an important role in controlling fat accumulation [21,22]. The *daf-2/INSR (lf*) causes fat accumulation in adults [23-25]. Food cues are sensed by an olfactory receptor in the amphidal sensory neurons and this, in turn, is relayed to the IIS pathway to control fat metabolism. However, the functions of INS on fat accumulation have not been identified. We studied the role of each of the forty INS on fat accumulation in adult worms and found that thirteen INS peptides (INS-3, 4, 6, 9, 11, 16, 18, 19, 25, 29, 30, 32 and DAF-28) had lower fat levels in comparison to wild type (Figure 5A and C). Seven INS peptides (INS-8, 12, 17, 21, 28, 37 and 39) elevated fat levels compared to wild type worms (Figure 5B). Our results suggest that most of the INS that act as agonists had reduced fat staining while most of the antagonists had increased fat staining. Of all the thirteen INS (*oe*) that could induce L1 arrest Q cell divisions (Figure 2D), most behaved as agonists for fat accumulation with the exception of INS-1, 2, 7 and 31 that had no effect, and INS-8 which acted as an antagonist for fat accumulation. Only INS-21 appeared to be specific for a role in fat accumulation exhibiting no effects on the other three phenotypes scored (Figure 5B).

**Figure 5.**
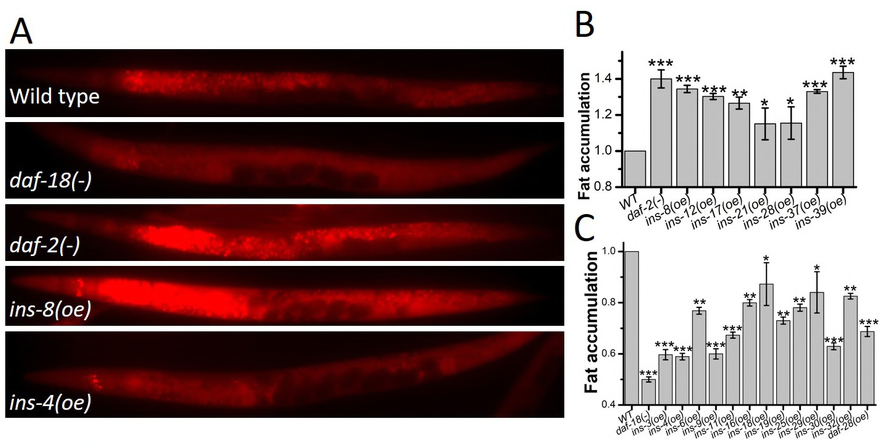
*ins* (*oe*) functions on fat accumulation. (A) Fat accumulation analyzed by nile red staining in wild type, *daf-18(-)*, *daf-2 (-)*, and *ins* (*oe*) worms. The *daf-2* and *daf-18* mutants have higher and lower level fat accumulation than wild type respectively. (B) 7 *ins* (*oe*) worms (antagonists) have higher fat accumulation than wild type. (C) 13 *ins* (*oe*) worms (agonists) have lower fat accumulation than wild type. Error bars represent the SD, * P value vs control <0.05, ** P value vs control <0.01 *** P value vs control <0.001. Also see details in Supplemental sheet.

### An F peptide and the β class INS act as agonists

According to our results, it is apparent that INS peptides that were structurally characterized as the β class and contain a sequence known as the F peptide are activators of the DAF-2/INSR(Figure 7). Nine of the β class INS peptides contain the F peptide (Figure 7). All of the β class INS behaved as agonists of DAF-2/INSR in our L1 arrest Q cell division assay, except for INS-10, which does not have an F peptide (Figure 7). We hypothesized that the F peptide contributes to the *C. elegans* INS activation in L1 arrest Q cell divisions. To test this hypothesis, we pan-neuronally expressed a strong agonist, INS-4 with a deletion of the F-peptide region or the F peptide alone and found that both failed to induce Q cell divisions during L1 arrest (Figure 6A and B). Pan-neuronal co-expression of the F peptide and the F peptide-lacking INS-4 resulted in the induction of Q cell divisions during L1 arrest (Figure 6A and B), showing that the F peptide can act in trans and is necessary for INS-4 activation of DAF-2/INSR. We then asked if co-expression of the F peptide with the β class INS-10, whose overexpression does not affect Q cell division, could induce L1 arrest Q cell divisions. The F peptide and INS-10 co-expression in trans failed to induce Q cell divisions during L1 arrest (Figure 6B). This suggests that the F peptide functionally complements INS-4 (minus F peptide) activity in a peptide sequence specific manner and/or that the pool of F peptides released upon processing of F peptide INS may not functionally complement INS peptides that lack an embedded F peptide sequence.

**Figure 6.**
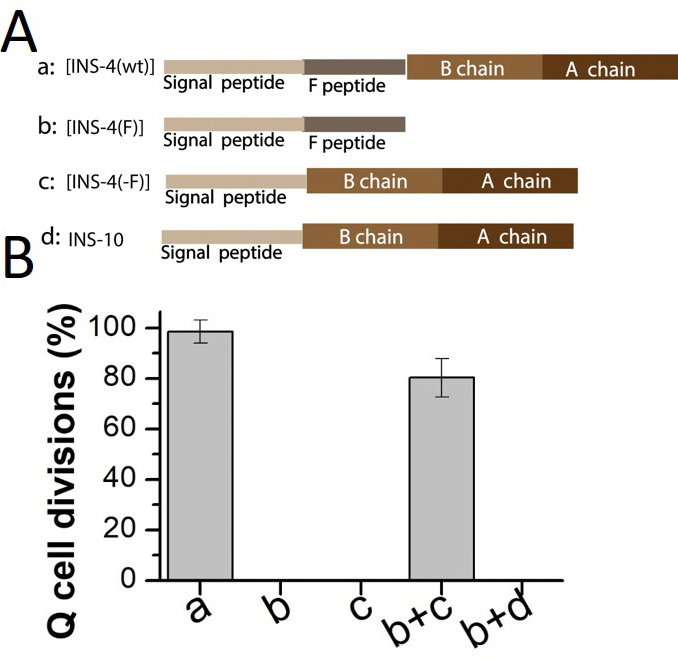
The F peptide is needed for INS-4 activation. (A) Variant of INS-4, a: wildtype *ins-4* [INS-4(wt)] b: *ins-4* F peptide only [INS-4(F)], c: *ins-4* with F peptide deleted [INS-4(-F)]. (B) INS-4 with no F peptide (c) or F peptide alone (b) does not induce L1 arrest Q cell divisions. However, adding back both in trans (b+c) can induce L1 arrest Q cell divisions. INS-10 has a structure similar to INS-4 (β class), but INS-10 has no F peptide. INS-10 + F peptide from INS-4 (b) co-injection does not induce L1 arrest Q cells divisions.

**Figure 7.**
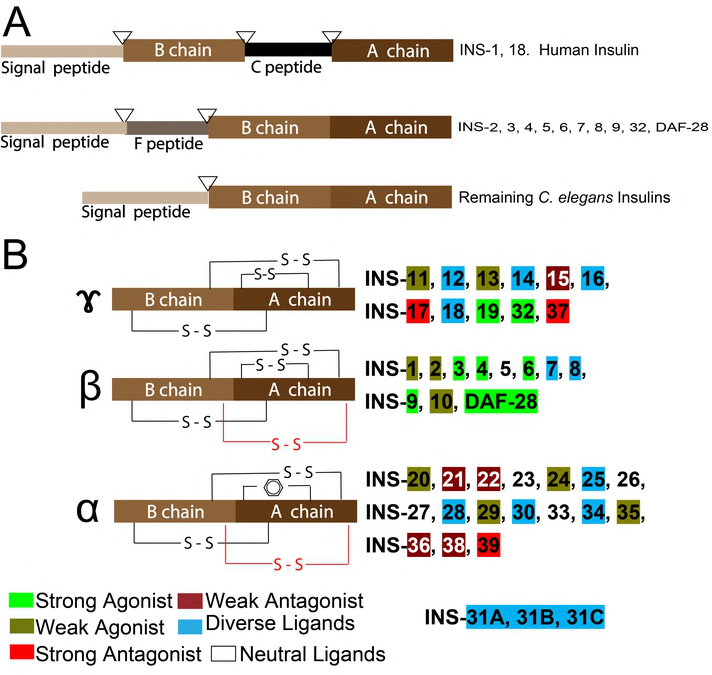
Insulin-like peptides in *C. elegans*. (A) All INS contain at least a signal peptide, B chain and C chain. Only INS-1 and INS-18 have a C peptide, like human insulin. F peptide is present in INS-2 through INS-9 and DAF-28. Predicated cleavage sites for the proteolytic processing (triangles). (B) *C. elegans* INS can be classified into three types based on disulfide bonds (PIERCE et al. 2001). Gamma insulins have the arrangement of three disulfide bonds as found in vertebrates while Alpha and Beta insulins contain an additional intra-chain disulfide bond (red). Alpha insulins lack the common intra-chain bond in the A chain, which is substituted by the interaction of aromatic amino acid side chains. INS-31 constitutes its own additional class with three repeats of B and A peptide chains. In our study we classified the 40 insulin ligands into 6 functional groups: strong agonist/antagonist: activity consistence within all tested phenotypes; weak agonist/antagonist: activity consistence within most tested phenotypes, but have no significant activity in other phenotypes; diverse: can have both agonist and antagonistic roles and neutral ligands: no significant activity in all tested IIS assays.

## Discussion

In this study, we created independent worm lines that overexpressed each of forty *C. elegans* INS peptides and assayed for IIS phenotypes in order to assign *in vivo* roles. The phenotypes that we measured in INS overexpression lines were dependent on processing by pro-protein convertase (EGL-3) in the nervous system. Although expressed in the nervous system, we showed that processed peptides must be secreted and can act cell non-autonomously on germline and M cells and is dependent on the DAF-2/INSR. Our study is the first to provide the functional data for all forty INS on L1 arrest life span, Q cell divisions, heat-stress induced dauer formation, and fat accumulation. Our results are summarized in Table 1 and Figure 7.

**Table 1.**
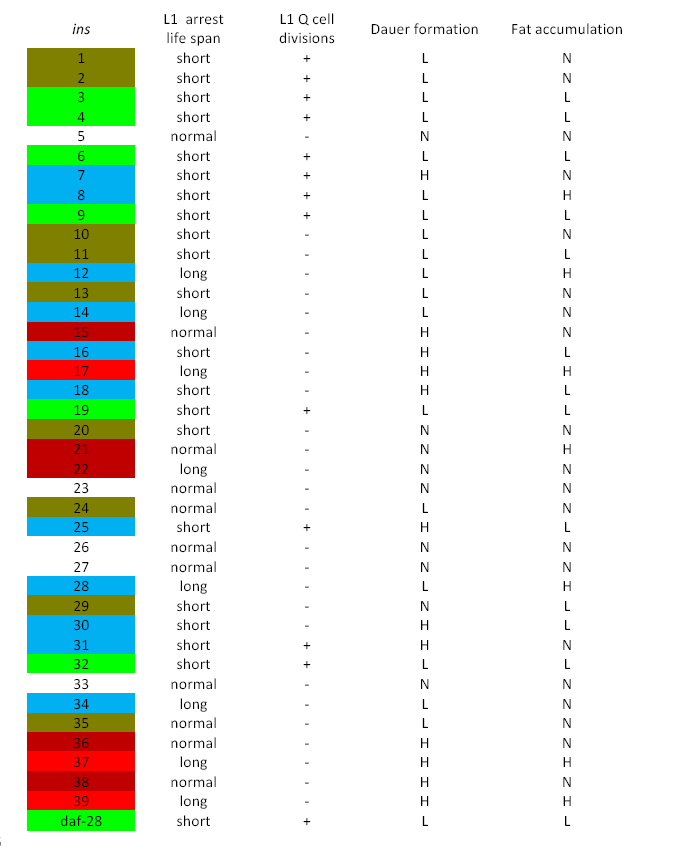
*C. elegans* INS peptides function data summary. Data are compared to wild type. +/-: with/without L1 arrest Q cell divisions. N: normal; L: low; H; high. See details in Supplemental sheet for raw data. Colour code: Strong agonist, Weak agonist, Strong antagonist, Weak antagonist, Diverse, Neutral.

| *ins* | L1 arrest<br>life span | L1 Q cell<br>divisions | Dauer formation | Fat accumulation |
| --- | --- | --- | --- | --- |
| 1 | short | + | L | N |
| 2 | short | + | L | N |
| 3 | short | + | L | L |
| 4 | short | + | L | L |
| 5 | normal | - | N | N |
| 6 | short | + | L | L |
| 7 | short | + | H | N |
| 8 | short | + | L | H |
| 9 | short | + | L | L |
| 10 | short | - | L | N |
| 11 | short | - | L | L |
| 12 | long | - | L | H |
| 13 | short | - | L | N |
| 14 | long | - | L | N |
| 15 | normal | - | H | N |
| 16 | short | - | H | L |
| 17 | long | - | H | H |
| 18 | short | - | H | L |
| 19 | short | + | L | L |
| 20 | short | - | N | N |
| 21 | normal | - | N | H |
| 22 | long | - | N | N |
| 23 | normal | - | N | N |
| 24 | normal | - | L | N |
| 25 | short | + | H | L |
| 26 | normal | - | N | N |
| 27 | normal | - | N | N |
| 28 | long | - | L | H |
| 29 | short | - | N | L |
| 30 | short | - | H | L |
| 31 | short | + | H | N |
| 32 | short | + | L | L |
| 33 | normal | - | N | N |
| 34 | long | - | L | N |
| 35 | normal | - | L | N |
| 36 | normal | - | H | N |
| 37 | long | - | H | H |
| 38 | normal | - | H | N |
| 39 | long | - | H | H |
| daf-28 | short | + | L | L |

Mutants with reduced IIS signaling have both a Daf-c phenotype and extended L1 arrest survival. We show that the INS-17, 37 and 39 are antagonists of DAF-2/INSR exhibiting increased L1 arrest survival and increased dauer formation. INS-17 was previously reported to work as a DAF-2/INSR antagonist for dauer regulation [11], but this work is the first to assign INS-37 and the 39 as antagonists. Our work also demonstrated that select INS peptides are DAF-2/INSR antagonists in controlling L1 arrest survival but are pleiotropic in their action in controlling dauer formation. For example, INS-12, 14, 28 and 34, which act as antagonists and can extend L1 arrest survival, can also act as agonists and have significantly lower dauer formation than wild type worms. On the other hand, INS-7, 16, 18, 25, 30, 31, which act as agonists in L1 arrest, but can have the opposite roles in dauer and act as an antagonist and increased dauer formation. Our results suggest that INS function in L1 arrested worms may be different from that of controlling dauer formation since it is an alternative L3 development stage, and thus INS peptides have spatiotemporal compartmentalization with respect to their function. Our finding is consistent with studies that show that dauer arrest and adult lifespan regulation by IIS are also decoupled [26-29].

Pan-neuronal INS overexpression that caused L1 arrest Q cell divisions identified thirteen INS peptides (INS-1, 2, 3, 4, 6, 7, 8, 9, 19, 25, 31, 32 and DAF-28) that act as agonists for the DAF-2/INSR. All thirteen INS peptides also have short L1 arrest life span, suggesting that all thirteen INS peptides behave as potential DAF-2/INSR agonists. Previous studies, based on differing assays assigned INS-3, 4, 6, 9, DAF-28 as potential agonist INS peptides [12,13]. These studies are consistent with our findings and support the reliability of our L1 arrest Q cell division readout as a means of categorizing the *ins* genes. INS-5 has been suggested to be an agonist [13], however, we did not find INS-5 to have any function in all of our assays, which is consistent with another report [12]. INS-1 was shown to be an antagonistic peptide based on dauer formation [7,12,30]. Overexpression of *ins-1*, enhances dauer arrest in weak *daf-2* mutants, suggesting that INS-1 antagonize DAF-2 insulin-like signaling. Also, INS-1 is antagonistic to DAF-2 for behavior [31]. However, in our assays we found INS-1 to have weak activation properties. INS-1 may be a complex peptide as INS-1 acts as an agonist for DAF-2 in salt chemotaxis learning [32].

We identified eight antagonistic INS peptides that could significantly extend L1 arrest life span and when overexpressed from the nervous system could suppress the *daf-18/pten* L1 arrest Q cell divisions. Thus, these INS peptides acted as therapeutic peptides for *daf-18/pten* worms. In humans, insulin and IGFs are thought to work as agonists and do not have antagonistic properties. Our work showed that *C. elegans* INS-6 is a strong agonist and INS-6 has been shown to bind and activate the human insulin receptor [33]. It would be interesting to know whether the antagonistic INS peptides we have identified in this study can bind to and inhibit the human insulin or IGF-1 receptor, if so, these *C. elegans* INS peptides could be used as future therapeutics.

Our study, revealed that INS-8 behaves as an agonist of DAF-2/INSR, because *ins-8 (oe)* shortens the lifespan of L1 arrested worms, has low penetrance dauer formation and promotes L1 arrest Q cell divisions. A previous study suggested that INS-8 may work as an agonist [8]. However, *ins-8 (oe)* worms behaved as an antagonist of DAF-2/INSR exhibiting higher fat accumulation. One study showed that *ins-8 (oe)* enhances *ins-7* mutant life span, which would suggest that INS-8 is an antagonist [8]. We suggest that the neuronal *ins-8* (*oe*) is sufficient to work as an agonist to activate the IIS pathway which in turn controls the L1 arrest life span and dauer formation, but in adult worms, it may work as an antagonist. This result with INS-8 is consistent with our finding that many INS peptides have distinct roles in mediating fat accumulation that is developmentally separate from its effects on dauer and L1 arrest life span. Insulin signaling temporally and in varying tissues of the body contributes differently to fat content [27].

Of the 40 INS peptides tested, eight appear to have specific functions in our phenotype assays. INS-15, INS-21 and INS-20, 22 act as DAF-2/INSR antagonists specifically for regulating dauer, fat metabolism, and L1 arrest life span respectively. Similarly, INS-24, 35, 36 and 38 only function in dauer formation.

To understand what makes an INS an activator we focused on the L1 arrest Q cell divisions as this assay determined with certainty which INS peptides acted as DAF-2/INSR activators. Our study revealed that the β class INS peptides which contains the three canonical disulfide bonds as well as an additional inter-chain disulfide bond are a good predictor of an INS peptide agonist. INS-1 to INS-10 and DAF-28 fall into this class (Figure 7). Nine of the β class INS contain an F peptide [7], the exception is INS-10. The F peptide is processed at the N-terminus by the signal peptidase cleavage site and at the C-terminus by either the proprotein convertase enzymes EGL-3/PC2-like with cleavage sequence (RR or KR) or a KPC-1/PC1-like site (R-X-X-R) [12] (Figure S1). INS-10 does have activation properties and reduces L1 arrest life span and dauer formation, but could not induce L1 arrest Q cell divisions. INS-5 was predicted to contain an F peptide [7], but upon further examination, INS-5 does not have a proprotein convertase site that would release the F peptide, but instead would be incorporated as the B chain (Figure 7, Figure S1). Thus, our results reveal a striking revelation that all INS peptides that are predicted to contain an F peptide should behave as agonists of the DAF-2/INSR (Figure 7). We showed that the F peptide is indeed required for INS-4 to induce L1 arrest Q cell divisions and the F peptide can be added back in trans to restore INS-4 (minus F peptide) function. Note that the F peptide is not an absolute requirement for an INS to induce L1 arrest Q cell divisions as INS-1, 19, 25, 31, and 32 could induce L1 arrest Q cell divisions (albeit not as strong as other eg. INS-4). Interestingly, the predicted signal sequences for INS-32 was longer than average INS peptides and therefore may produce an F peptide. This prompted us to look more closely at the predicted peptides and using the SignalP 4.1 (http://www.cbs.dtu.dk/services/SignalP/), we identified INS-32 as having potential F peptide (Figure S1). In addition, we also propose that INS-19 has an F peptide as it has a potential pro-protein convertase cleavage site (Figure S1).

Human insulin has a C peptide, and human IGF-1 and IGF-2 have E peptides that are cleaved during processing analogous to the F peptides identified in *C. elegans* INS peptides. Our work on the F peptide should stimulate closer examination of peptides released upon processing of human Insulin and IGF. For instance, IGF-1 is one of the key molecules in cancer biology, however little is known about the role of the E peptide. E peptide is thought to have functional properties as the release from IGF-1 thought to induce cellular proliferation in the human prostate cancer [34]. The C peptide of proinsulin is important in processing of mature insulin and may have biological activity as a report suggests that it binds to a G protein-coupled surface receptor and activates Ca (2+)-dependent intracellular signaling pathways [35]. Since we have provided evidence in *C. elegans* that the F peptide can work in trans with the INS-4 lacking an F peptide, the F peptide serves as a modulator of INS-4 to induce L1 arrest Q cell divisions.

Finally, of the 40 INS (*oe*) strains tested, INS-5, 23, 26, 27, 33 were not functional in the selected assays. These INS peptides may have specific roles that have not been uncovered through the assays selected. The INS peptides are also thought to function in a combinatorial fashion and perhaps these single INS peptides have no function on their own and may participate with the other INS to exert their function [14]. Alternatively, these INS may bind to receptors other than DAF-2/INSR. A report has suggested that additional insulin-like receptors have been identified in the *C. elegans* genome [36].

In conclusion, our work systematically tested the functions of each of forty INS on dauer formation, L1 arrest life span, L1 arrest Q cell divisions and fat accumulation phenotypes (Table 1). By using these IIS phenotypes as readouts of insulin peptide activity, we found that seven INS peptides (3, 4, 6, 9, 19, 32 and DAF-28) were strong agonists and three INS peptides (17, 37 and 39) were strong antagonists of DAF-2/INSR, because these INS peptides acted either as agonists or antagonists in all our tested phenotypes. Five INS peptides (15, 21, 22, 36, and 38) were found to be weak antagonists; and nine INS peptides (1, 2, 10, 11, 13, 20, 24, 29 and 35) were weak agonists. Five INS peptides (5, 23, 26, 27 and 33) were neutral ligands. Eleven INS peptides (INS-7, 8, 12, 14, 16, 18, 25, 28, 30, 31 and 34) have different roles in different stress environments and developmental stages (Table 1 and Figure 7). These diverse functions of INS may contribute to important influences on development, metabolism, and aging-related diseases.

## Methods

### Strains

Most of the strains used in this study were acquired from the *Caenorhabditis* Genetics Center (CGC). Standard culture methods were used as previously described [37]. Strains were grown on OP50 *E. coli* and cultured at 20°C unless otherwise indicated. Strains used in this study: CZ10175: *zdIs5 [mec-4p∷GFP + lin-15(+)] I*, RB712: *daf-18(ok480)*, CB1370: *daf-2 (e1370)*, DR1942: *daf-2*(*e979*), VC671: *egl-3 (ok979)*. PD4666: *ayIs6 [Phlh-8∷gfp + dpy-20(+)].*

### Transgenic strains

For the *ins* overexpression strains, the insulin genomic sequences were amplified from a N2 genomic DNA and were placed under control of the pan neuronal promoter P*rgef-1* by standard cloning procedures [15]. A plasmid with the injection marker *odr-1∷rfp* was injected into *Pmec-4∷GFP(zdIs5)* [38] worms using standard microinjection methods [39]. For the F peptide experiment: Q5 mutagenesis (NEB) using primers was used to delete the F-peptide sequence from the *Prgef-1∷*INS-4 plasmid. Each injected strain had at least 2 stable lines.

### L1 arrest Q Cell divisions

Non-starved well-maintained mix staged worms were collected to prepare embryos, as described [40]. In brief, embryos were maintained and hatched in sterile M9 and incubated at 20°C with low speed rocking to initiate L1 arrest. The final Q cell descendants (A/PVM) were observed under an Axioplan fluorescent microscope (Zeiss, Germany) after 2 days or more in L1 arrest. 50-100 µL of M9 containing greater than 50 L1 arrested worms were removed from the culture. The total number of worms and the worms with A/PVM cells divisions were counted. For transgenic strains, only the worms with the injection marker were counted and analyzed.

Similarly, M cell divisions were analyzed by using *ayIs6* strains [41].

### Antibody staining

Antibody staining was performed as previously described (Chin-Sang et al. 1999). To detect germline cells, rabbit anti-PGL-1 (P-granule component) (1:20000) (a gift from Dr. Susan Strome) was used as the primary antibody. Detection was with a FITC-labeled goat anti-rabbit secondary antibody (1:100). For transgenic strains, only the worms with the injection marker were counted and analyzed. The total number of worms and the worms with germ-cell divisions were counted. Analysis of worms was using an Axioplan fluorescent microscope (Zeiss, Germany).

### L1 arrest life span assays

Life span was assessed in liquid medium [18]. L1 worms were cultured in 1 mL M9, 50-100 µL was taken to ensure the sample size was larger than 50, and the worms were scored every day. We scored survival by counting the number of worms that were moving (alive) and then dividing that number by the total number of worms in the aliquot. To compare the survival rates between strains, the L1 arrests were carried out in triplicate with at least 100 L1s and the mean survival rate calculated by the Kaplan-Meier method [42], that is the fraction of living animals over a time course. The significance of difference in overall survival rate is performed using the log-rank test [43].

### Fat staining

Synchronized eggs were cultured on OP50 plates with 25 ng/ml of Nile red for 3 days at 20°C, and then washed 3 times with M9, cultured on normal OP50 plates for 1 more day at 20°C to eliminate the Nile red OP50 background in the intestine. Worms were collected and washed in M9 3 times, then fixed in 40% isopropanol for 3 min. At least 30 animals were imaged in at least three separate experiments using a Zeiss Axioplan. The fluorescent intensity in the whole worm was quantified by using ImageJ.

### Dauer formation at high temperature

We analyzed L2 dauer formation at 29°C as synchronized zdIs5 worm eggs hatched at 29°C presented a higher percentage dauer phenotype. The dauer, dauer-like and adult worms with injection marker were counted. The dauer percentages were calculated. Three independent trails were performed for each strain, each sample size was greater than 50.

## Acknowledgments

We are grateful to *Caenorhabditis* Genomic Center for providing strains, which is funded by the NIH Office of Research Infrastructure Programs (P40OD010440). The work is supported by grants from the Natural Sciences and Engineering Research Council of Canada (NSERC 249779) and the Canadian Institutes of Health Research (CIHR 130541).

## Supporting Information Legends

**File: S1_Peptide sequences and Strain names**

Figure S1: Predicted peptides of the *C. elegans* Insulin Like Peptides (INS)

Predicted and revised peptide sequences of the 40 *C. elegans* INS. The signal sequence peptide, B peptide, A peptide, C peptide and F peptide are colour coded as indicated. INS-19 and INS-32 are revised based on our work.

Table S1. INS strains

INS overexpressing strains used in this study. Strain names and alleles are indicated.

**File: S2_raw data**

An Excel workbook with the raw data for the 4 phenotypes scored in this study.

